# Distinct subcellular localization of a Type I CRISPR complex and the Cas3 nuclease in bacteria

**DOI:** 10.1101/2020.09.29.318501

**Authors:** Sutharsan Govindarajan, Adair Borges, Joseph Bondy-Denomy

## Abstract

CRISPR-Cas systems are prokaryotic adaptive immune systems that have been well characterized biochemically, but *in vivo* spatiotemporal regulation and cell biology remains largely unaddressed. Here, we used fluorescent fusion proteins to study the localization of the Type I-F CRISPR-Cas system native to *Pseudomonas aeruginosa*. When targeted to an integrated prophage, the crRNA-guided (Csy) complex and a majority of Cas3 molecules in the cell are recruited to a single focus. When lacking a target in the cell, however, the Csy complex is broadly nucleoid bound, while Cas3 is diffuse in the cytoplasm. Nucleoid association for the Csy proteins is crRNA-dependent, and inhibited by expression of anti-CRISPR AcrIF2, which blocks PAM binding. The Cas9 nuclease is also nucleoid localized, only when gRNA-bound, which is abolished by PAM mimic, AcrIIA4. Our findings reveal PAM-dependent nucleoid surveillance and spatiotemporal regulation in Type I CRISPR-Cas that separates the nuclease-helicase Cas3 from the crRNA-guided surveillance complex.

## Introduction

Bacteria have evolved a wide range of immune mechanisms, including the clustered regularly interspaced short palindromic repeats (CRISPR) and CRISPR-associated (cas) genes, to protect from bacteriophages and other mobile genetic elements (Marraffini, 2015). The adaptive CRISPR-Cas system is present in almost 85% of archaea and 40% of bacterial genomes sequenced. Currently, CRISPR-Cas systems are categorized into 2 broad classes, 6 types and 33 subtypes (Makarova *et al*., 2019). CRISPR-Cas systems acquire foreign DNA into the CRISPR array as new ‘spacers’, transcribes and processes that array to generate CRISPR RNAs (crRNAs), which complex with Cas proteins. This crRNA-guided complex surveils the cell for a protospacer adjacent motif (PAM), and subsequent complementary base pairing with the crRNA, which triggers cleavage of the invading nucleic acid. In the case of type I CRISPR-Cas systems, which are the most abundant in bacteria (Van Der Oost *et al*., 2014; Makarova *et al*., 2019), a multi-subunit crRNA-guided surveillance complex (Cascade in Type I-E, Csy complex in Type I-F) recognizes the PAM and protospacer, and then recruits a *trans*-acting helicase-nuclease (Cas3) for target degradation (Redding *et al*., 2015).

How CRISPR-Cas effectors are organized within cells is currently not well understood. For efficiently functioning as a defense system, CRISPR-Cas must rapidly recognize the incoming foreign DNA and destroy it before it becomes established. Previous *in vitro* studies using single molecule imaging has shown that Type I-E Cascade (which is the crRNA-guided complex of the I-E system) spends between 0.1 and 10 s scanning targets in DNA (Redding *et al*., 2015; Xue *et al*., 2017; Dillard *et al*., 2018). A recent *in vivo* study using super resolution microscopy and single molecule tracking of cascade in live *E. coli* suggested a timescale of 30 ms for target probing (Vink *et al*., 2020), and similar binding kinetics have also been suggested for Cas9 (Jones *et al*., 2017; Martens *et al*., 2019). Cascade was suggested to spend approximately 50% of its search time on DNA and the rest distributed in the cytoplasm, however, the impact of phage infection and Cas3 localization have not determined. Additionally, ChIP-seq analysis of Cascade-DNA interactions in *E. coli* revealed that less than 5 bp of crRNA-DNA interaction is sufficient to promote association of cascade at numerous sites in the genome (Cooper *et al*., 2018). Genomic associations of CRISPR-Cas stands in contrast to another important class of defense systems, restriction modification (R-M). Type I R–M complexes (HsdRMS) localize to the inner membrane in such a way that their activities are controlled spatially. The methyltransferase is exposed in the cytoplasmic-side of the inner membrane, for accessing the host DNA, while the restriction enzyme components are positioned in the periplasmic-side of the membrane (Holubová *et al*., 2000; Holubova *et al*., 2004).

The type I-F CRISPR-Cas system of *P. aeruginosa* has emerged as a powerful model for understanding various aspects of CRISPR-Cas biology, including a mechanistic understanding of type I systems (Wiedenheft *et al*., 2011; Guo *et al*., 2017; Rollins *et al*., 2019), the discovery (Bondy-Denomy *et al*., 2013) and *in vivo* characterization (Borges *et al*., 2018; Landsberger *et al*., 2018) of phage-encoded anti-CRISPR proteins (Acrs), and identification of regulatory pathways governing CRISPR-Cas expression (Høyland-Kroghsbo *et al*., 2017; Lin *et al*., 2019a; Lin *et al*., 2019b; Ahator *et al*., 2020; Borges *et al*., 2020). The I-F system of *P. aeruginosa* PA14 consists of two CRISPR loci and six Cas proteins, namely Csy1-4, which form the Csy complex (surveillance complex of the I-F system), Cas3 (a fusion of Cas2-3), a trans-acting nuclease/helicase protein, and Cas1, which drives spacer acquisition (Wiedenheft *et al*., 2011; Rollins *et al*., 2017). Structural and biochemical studies have shown that Csy1-4 assemble on a 60 nucleotide crRNA to form a 350 kDa sea-horse-shaped crRNA-guided surveillance complex (Chowdhury *et al*., 2017; Guo *et al*., 2017; Rollins *et al*., 2019). The Csy surveillance complex recognizes DNA first via Csy1-PAM interaction (G-G/C-C), leading to destabilization of the DNA duplex, strand invasion, R-loop formation in the seed region, and downstream base pairing. Finally, DNA-bound Csy complex triggers recruitment of Cas3 nuclease, which mediates processive degradation of the target DNA in a 3’–5’ direction (Chowdhury *et al*., 2017; Guo *et al*., 2017; Rollins *et al*., 2019).

Here, we address the cell biology of a naturally active *P. aeruginosa* Type I-F CRISPR-Cas system by directly observing its subcellular localization using live cell microscopy. Using functional fluorescent fusions chromosomally integrated at the native locus, we show that the Csy1 and Csy4 proteins, which are part of the surveillance complex, are largely nucleoid bound, while Cas3 nuclease is cytoplasmic, but both can be re-directed to a stable intracellular target (i.e. a prophage). When the Csy complex is formed with a crRNA, it binds the nucleoid even in the absence of a target, but the individual Cas proteins do not. Nucleoid localization of the Csy complex, and Cas9, is mediated at the level of PAM recognition and is specifically disrupted by PAM-mimetic anti-CRISPR proteins. Taken together, our study suggests that the relatively promiscuous PAM-dependent (e.g. 5’-GG-3’) DNA search mechanism likely relegates these complexes to predominantly associate with the host genome, without an active mechanism for preventing host genome surveillance.

## Results

### The majority of endogenous Cas3 molecules are recruited to CRISPR targets

To investigate CRISPR-Cas subcellular localization, we focused on *P. aeruginosa* strain UCBPP-PA14 (denoted PA14), which has a type I-F CRISPR-Cas system (Cady *et al*., 2012). We constructed PA14 strains in which Csy1 (Cas8), Csy4 (Cas6) and Cas3 are fused with sfCherry at their native locus. We assessed the functionality of the tagged strains using a panel of isogenic phages to read out CRISPR-Cas function: DMS3 (untargeted control), DMS3m (targeted by a natural spacer, CRISPR2 spacer 1), and DMS3m engineered phages that expresses Acr proteins AcrIF1, AcrIF2, AcrIF3 or AcrIF4 (Borges *et al*., 2018). All three fusions exhibited CRISPR-Cas activity similar to the wild-type and were inhibited by Acr proteins (Fig. S1). Using these strains as reporters, we performed live cell fluorescence microscopy to visualize the distribution of the Cas proteins. We observed that all three proteins appeared diffuse in the cytoplasm under this condition (Fig. 1a). Of note, cells had to be grown to high cell density, which is important for Cas protein expression in *P. aeruginosa* (Fig. 1b) (Høyland-Kroghsbo *et al*., 2017; Borges *et al*., 2020). Analysis of fluorescent signal from single cells indicated that Csy1 and Csy4 protein levels are higher than Cas3 (Fig. 1c).

**Figure 1.**
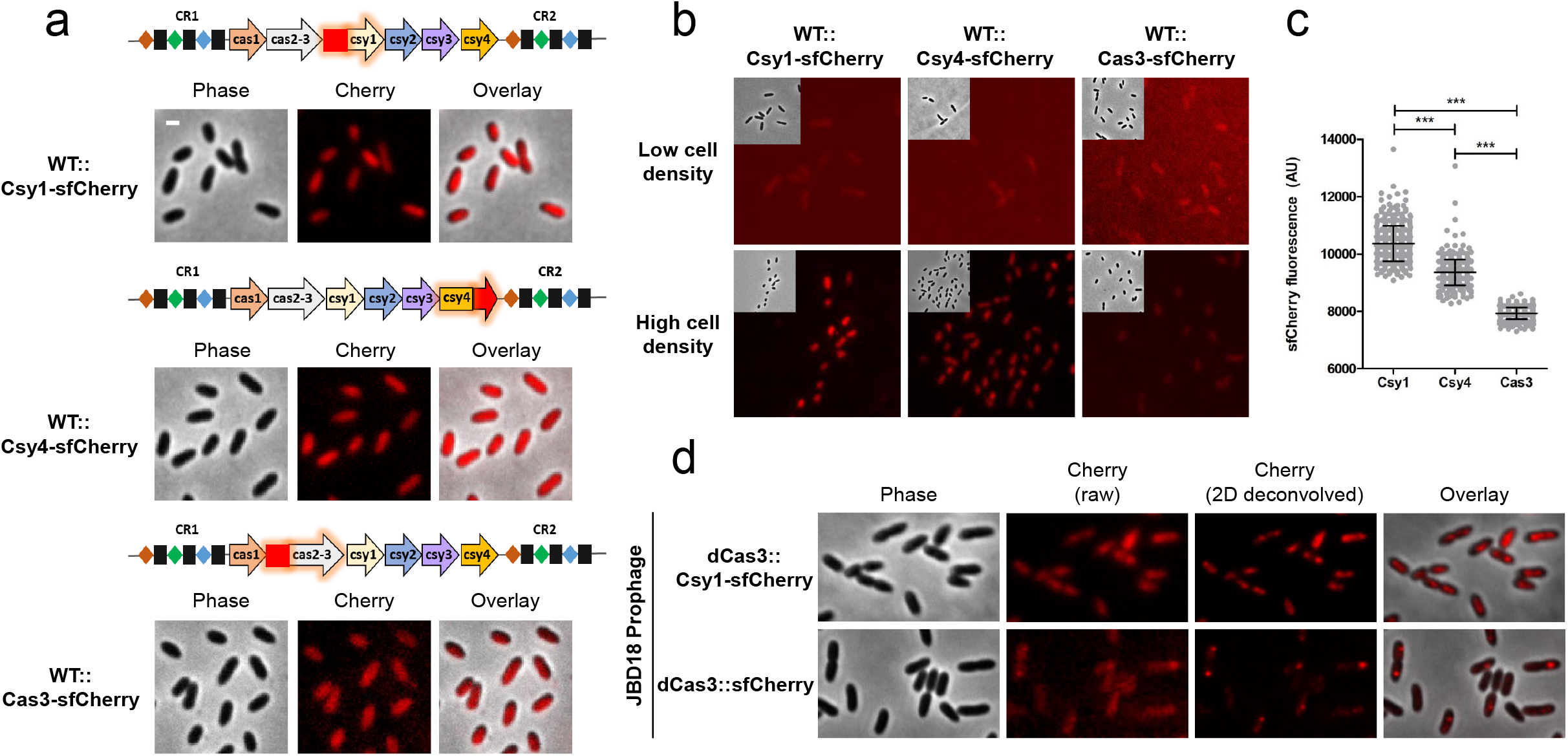
Live-cell fluorescence imaging of *P. aeruginosa* PA14 expressing type I-F Cas protein reporters from endogenous locus. (a) Fluorescence microscopy of wild-type PA14 expressing Csy1-sfCherry, Csy4-sfCherry, or Cas3-sfCherry. (b) Comparison of fluorescence in cells expressing Csy1-sfCherry, Csy4-sfCherry, or Cas3-sfCherry at low cell density (OD600 = 0.2 - 0.3) or high cell density (OD600 = 1 - 1.5). (c) sfCherry fluorescence (AU) in wild-type cells grown to high cell density expressing sfCherry-tagged Csy1, Csy4, or Cas3. More than 200 cells from each genotype were analyzed from two independent experiments. Means and standard deviations are shown. The statistical significance was calculated using unpaired t-test analysis (*** p<0.0001). (d) Fluorescence microscopy of PA14 expressing Csy1-sfCherry (in dCas3 background) or dCas3-sfCherry with a JBD18 prophage. sfCherry fluorescence shown in raw as well as deconvolved images. Scale bar, 1 μm.

To observe fluorescent Cas proteins localizing to target DNA *in vivo*, we generated lysogenic strains containing JBD18 as a prophage in PA14 strains expressing chromosomal Csy1-sfCherry (in dCas3 background) or dCas3-sfCherry. JBD18 naturally has five protospacer sequences with the correct PAM (Cady *et al*., 2012). Both Csy1-sfCherry and dCas3-sfCherry formed fluorescent foci in the presence of the JBD18 prophage (Fig. 1d). Notably, the foci formed by dCas3-sfCherry were quite discrete, suggesting that most of the cellular dCas3 protein is recruited to the targeted locus. Taken together, the data presented above shows that endogenous levels of tagged I-F Cas proteins are functional, can be visualized, and can read out *bona fide* target DNA binding events *in vivo*, which apparently recruits multiple Cas3 proteins.

### Csy complex is nucleoid-enriched while the Cas3 nuclease is cytoplasmic

To determine the localization pattern of the Csy complex, we took an approach that can clearly differentiate membrane associated, DNA bound and cytoplasmic proteins (Hershko-Shalev *et al*., 2016). Cells were treated with the DNA-damaging antibiotic nalidixic acid (NA). At the concentration used, *P. aeruginosa* continues to grow slowly and forms long cells with compacted nucleoid (schematic presented in Fig. 2a). In these nucleoid compacted cells, DNA-localized proteins can be clearly differentiated from cytoplasmic and membrane proteins. We observed that Csy1-sfCherry and Csy4-sfCherry are enriched in the nucleoid, as evidenced by their colocalization with DAPI stain, while Cas3-sfCherry appeared diffuse in the cytoplasm (Fig. 2b). To confirm that nucleoid localization of Csy1 and Csy4 did not occur as a result of DNA damage caused by NA treatment, we treated cells with chloramphenicol (200 μg/ml), which causes nucleoid compaction as a result of translation inhibition, leading to transertion inhibition (Van Helvoort *et al*., 1996; Govindarajan *et al*., 2013). Also, in these nucleoid compacted cells, Csy1 and Csy4, but not Cas3, were found to be nucleoid-enriched (Fig. S2).

**Figure 2.**
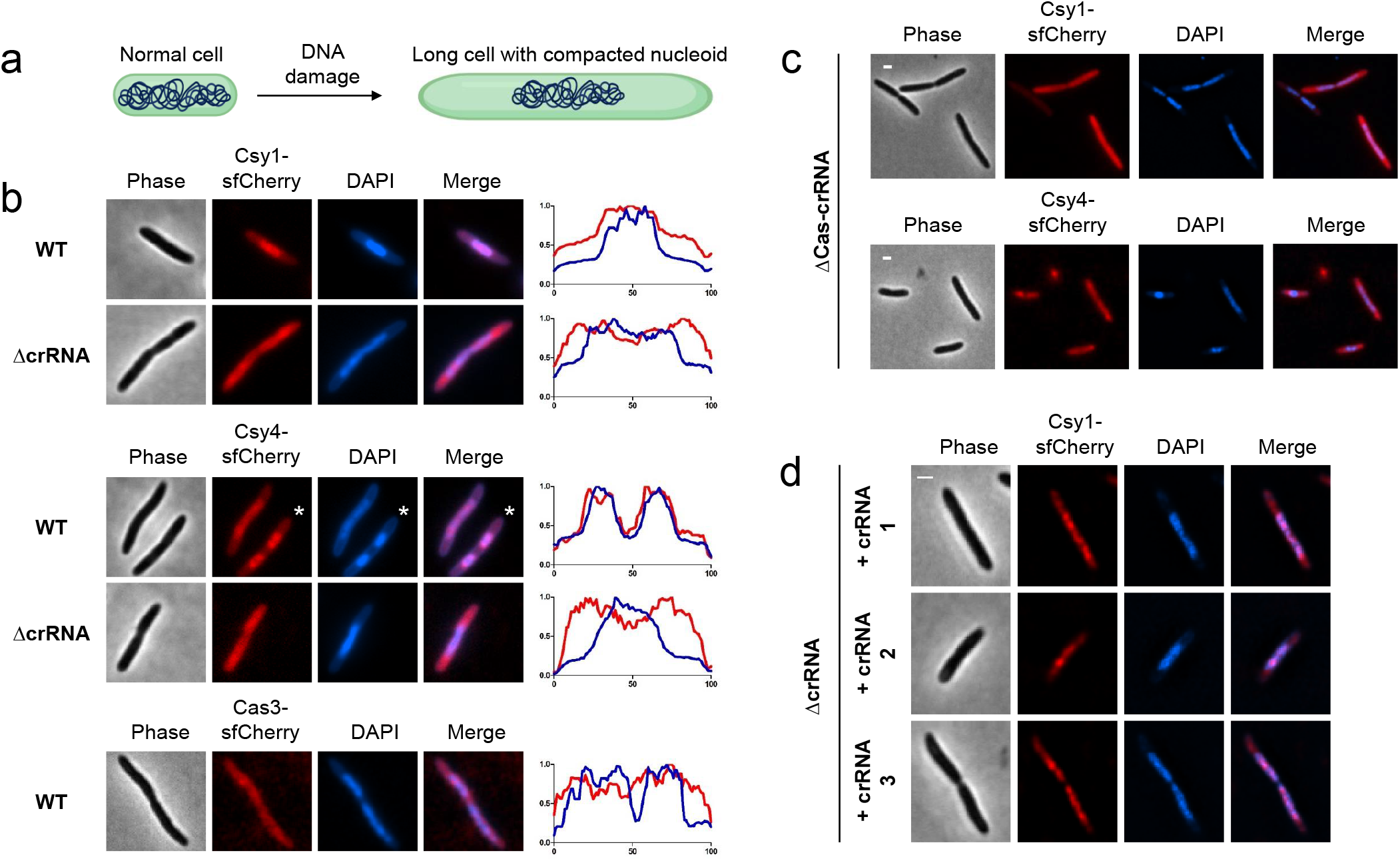
Csy complex is nucleoid-enriched in a crRNA-dependent manner while Cas3 is cytoplasmic. (a) A model for DNA damage induced nucleoid compaction. (b) Fluorescence microscopy of nucleoid compacted wild-type or ΔcrRNA cells expressing Csy1-sfCherry or Csy4-sfCherry and nucleoid compacted wild-type cells expressing Cas3-sfCherry. Fluorescence intensity map plotted against the long cell axis are shown. * indicates which cell is plotted in the map. (c) Fluorescence microscopy of nucleoid compacted ΔCas-crRNA cells expressing Csy1-sfCherry or Csy4-sfCherry from a plasmid. (d) Fluorescence microscopy of nucleoid compacted ΔcrRNA cells expressing Csy1-sfCherry and synthetic crRNAs. Scale bars, 1 μm.

We next asked whether Csy protein nucleoid localization is dependent on crRNA-mediated surveillance complex formation. To test this, the same Csy1-sfCherry and Csy4-sfCherry labels were inserted into the chromosome of a ΔCRISPR strain that lacks all 35 spacers, but still expresses all Cas proteins. Both Csy1-sfCherry and Csy4-sfCherry lost nucleoid localization in the ΔCRISPR mutant (Fig. 2b) suggesting that crRNA-mediated assembly of the surveillance complex is essential for nucleoid localization. Additionally, expression of Csy1-sfCherry or Csy4-sfCherry ectopically from a plasmid in NA-treated cells that lack all components of the CRISPR-Cas system (ΔCRISPR-Cas), showed that they were not enriched in the nucleoid (Fig. 2c).

PA14 has a total of 35 distinct crRNAs produced from its two CRISPR loci. It is possible that one or more of these crRNAs with partial match to the genomic DNA is responsible for the surveillance complex nucleoid localization. To determine the importance of crRNAs in nucleoid surveillance, we expressed synthetic crRNAs, which do not have a detectable sequence match in the PA14 genome, in ΔCRISPR cells and asked if they can restore Csy1-sfCherry nucleoid localization. All three crRNAs restored Csy1-sfCherry nucleoid localization in ΔCRISPR cells suggesting that the Csy complex, once assembled, is intrinsically capable of binding the nucleoid independent of the crRNA sequence (Fig. 2d). Together, these data indicate that nucleoid-localization of Csy1 and Csy4 is not an intrinsic property of free protein, but occurs as a result of functional surveillance complex formation, while the Cas3 nuclease is spread throughout the cell.

### An Anti-CRISPR that blocks PAM binding specifically prevents nucleoid localization

Next, we wanted to determine the mechanism of nucleoid binding and understand whether the surveillance complex binds to the nucleoid directly or through its interaction with other host factors. The surveillance complex recognizes target DNA in two steps: (i) interaction with the ‘GG’ PAM, which is mediated mostly by residues in Csy1 and (ii) base pairing between the crRNA with target DNA, which is mediated by the spacer region of the crRNA (Chowdhury *et al*., 2017; Guo *et al*., 2017; Rollins *et al*., 2019). Two Type I-F Acr proteins have been identified that distinguish between these two binding mechanisms (Bondy-Denomy*et al*., 2013; Bondy-Denomy *et al*., 2015; Maxwell *et al*., 2016; Chowdhury *et al*., 2017; Guo *et al*., 2017). AcrIF1 binds to Csy3, blocking crRNA-DNA target hybridization but does not occlude the PAM binding site, while AcrIF2 binds to the Csy1-Csy2 heterodimer, and specifically competes with PAM binding in target DNA. AcrIF3 was also utilized, which binds to Cas3 nuclease and prevents target cleavage but does not have any known effect on surveillance complex (Bondy-Denomy *et al*., 2015; Wang *et al*., 2016). Csy1-sfCherry localization was monitored in NA-treated nucleoid compacted cells expressing one of these three anti-CRISPRs, either from a plasmid (Fig. 3a) or from isogenic DMS3m prophages (Borges *et al*., 2018)(Fig. 3b) integrated in the reporter strain. Csy1-sfCherry localization to nucleoid was not affected when AcrIF1 or AcrIF3 were expressed. However, in cells expressing AcrIF2, nucleoid localization of Csy1-sfCherry is completely disrupted (Fig. 3a and 3b). Of note, prophage containing cells were transformed with a plasmid expressing the C-repressor in order to prevent possible excision during NA treatment. Finally, the expression of recently characterized AcrIF11 (Marino *et al*., 2018; Niu *et al*., 2020), an enzyme that ADP-ribosylates Csy1 to prevent PAM binding, also disrupted nucleoid localization (Fig. S3). Together, these results suggest that PAM recognition is the dominant and direct factor that mediates Csy complex localization to nucleoid.

**Figure 3.**
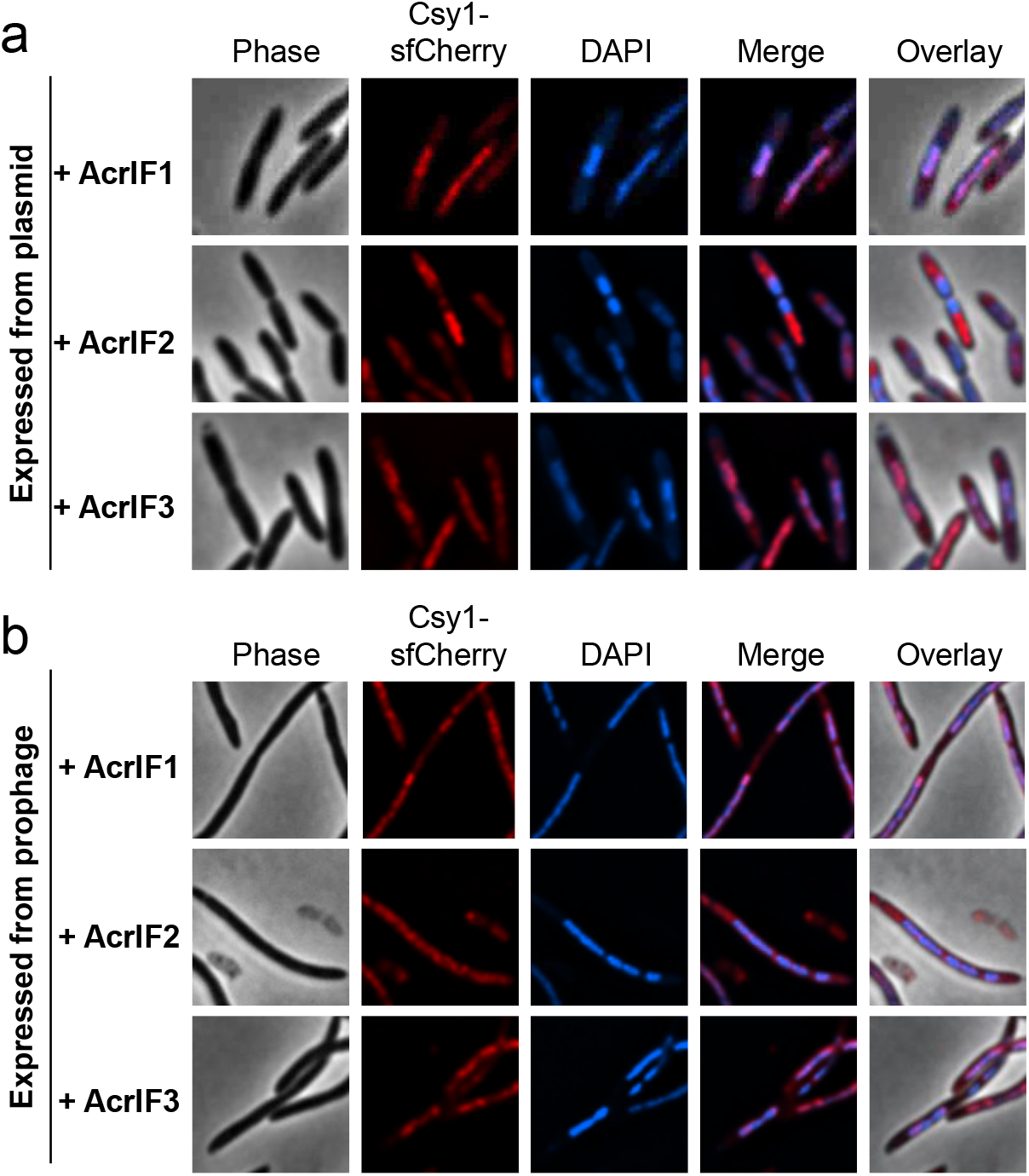
AcrIF2 but not AcrIF1 or AcrIF3 blocks nucleoid localization of the Csy complex. Fluorescence microscopy of nucleoid compacted wild-type cells producing Csy1-sfCherry and expressing anti-CRISPR proteins AcrIF1, AcrIF2 or AcrIF3 from (a) a plasmid or (b) a prophage.

### Phage infection does not modulate Csy complex localization

As a defense system against phage, we next wanted to address the important question of whether CRISPR-Cas distribution responds to phage infection. To enable visualization of infecting phage particles, we propagated DMS3 (0 protospacers) and DMS3m (1 protospacer) in a strain that expressed GpT, the major capsid protein, fused with mNeonGreen fluorescent protein. This allowed us to obtain fluorescent mosaic phage particles that contain labelled as well as unlabeled GpT protein. We verified that 100% of fluorescent phages packed viral DNA, as evidenced by colocalization of mNeonGreen and DAPI stain (Fig. 4a). The subcellular localization of Csy1-sfCherry and Cas3-sfCherry were assessed in the presence of labelled DMS3 (untargeted phage) or DMS3m (targeted phage) after 15 minutes of infection. In both cases, the apparent diffuse localization of Csy1-sfCherry was not affected at a gross level, when comparing cells with fluorescent phage particles adjacent to the cell surface to those without (Fig. 4b and 4c). To check whether presence of more protospacers would enable recruitment of Csy complex or Cas3, we infected Csy1-sfCherry (in dCas3 background) and dCas3-sfCherry expressing cells with unlabeled JBD18 under the same conditions (Fig. S4). Also, here we did not observe any change in localization despite this being the same genotype where JBD18 prophage recognition was so striking (Fig. 1d). These data suggest that phage infection, irrespective of presence or absence of a target sequence, does not significantly modulate the distribution of the Csy complex or Cas3.

**Figure 4.**
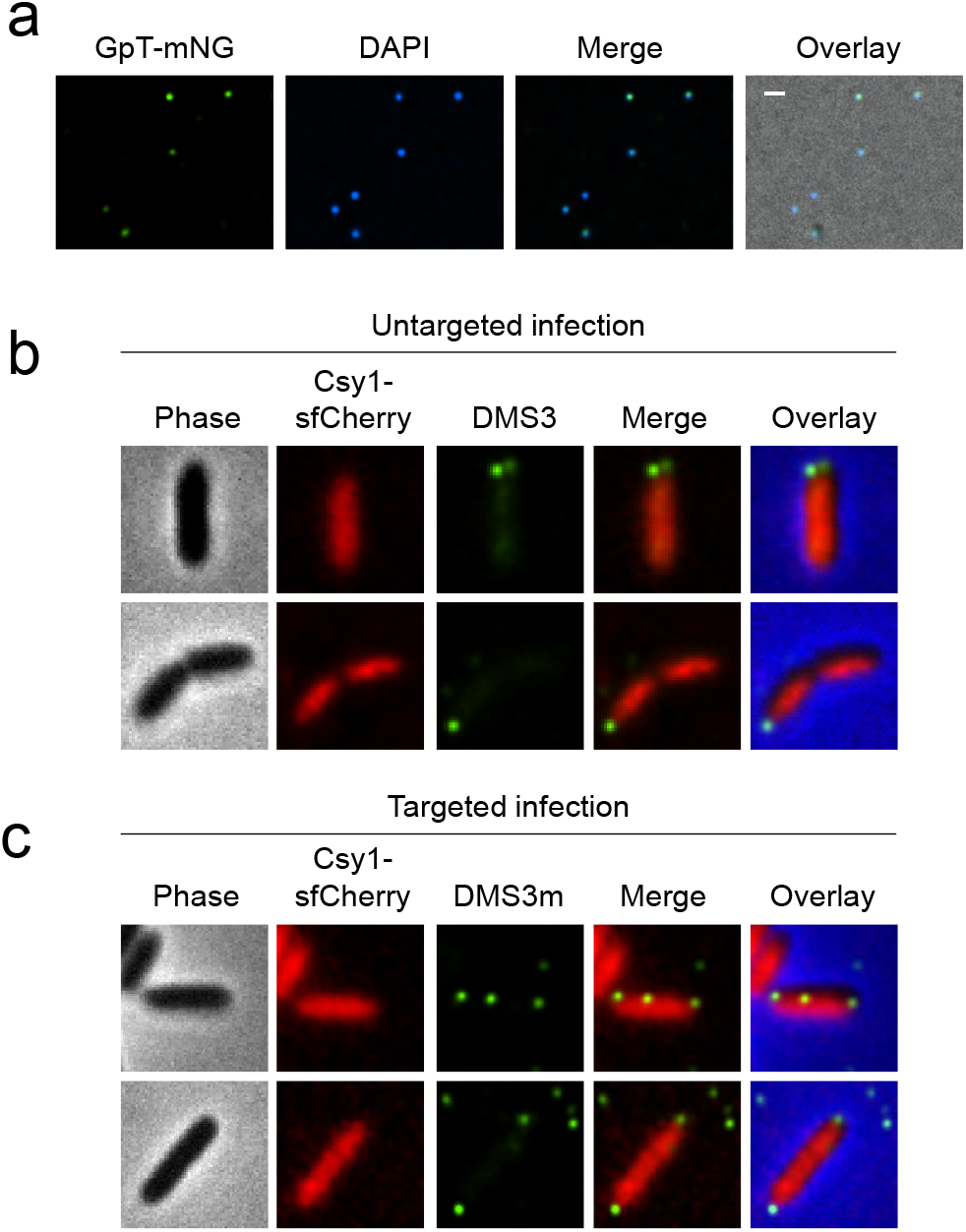
Phage infection does not affect Csy localization. (a) Fluorescence microscopy images of DMS3 phages with capsid labelled with GpT-mNeonGreen and DNA labelled with DAPI. Fluorescence microscopy of Csy1-sfCherry producing cells (in dCas3 background) infected with labelled phages (b) DMS3 (0 protospacers) or (c) or DMS3m (1 protospacer). Scale bar, 1 μm.

### PAM-dependent nucleoid localization is conserved for Spy Cas9

Having observed that the multi-subunit type I-F Csy surveillance complex is nucleoid enriched, we next wondered whether nucleoid localization is a conserved property of Class 2 CRISPR-Cas single protein effectors. For this purpose, we chose Cas9 of *Streptococcus pyogenes* as a model protein, which coincidentally uses the same PAM as the Csy complex. To test the localization of SpyCas9, we used a plasmid that expressed a functional SpyCas9 fused with cherry (Mendoza *et al*., 2020) in *P. aeruginosa* PA01 and monitored its localization in the presence and absence of a crRNA in NA-treated nucleoid compacted cells. Interestingly, SpyCas9 was found to be diffuse in its Apo form (i.e. lacking a crRNA) and nucleoid localized when complexed with a crRNA, provided as a single guide RNA (Fig. 5). When we expressed AcrIIA4, an anti-CRISPR that inhibits SpyCas9 by competing with the PAM-interacting domain (Dong *et al*., 2017; Rauch *et al*., 2017), nucleoid localization of guide RNA bound SpyCas9 was abolished (Fig. 5). These data, taken together with previous reports (Jones *et al*., 2017; Martens *et al*., 2019; Vink *et al*., 2020), suggests that genome surveillance via PAM-dependent nucleoid localization appears to be a conserved property of type I and type II CRISPR-Cas systems.

**Figure 5.**
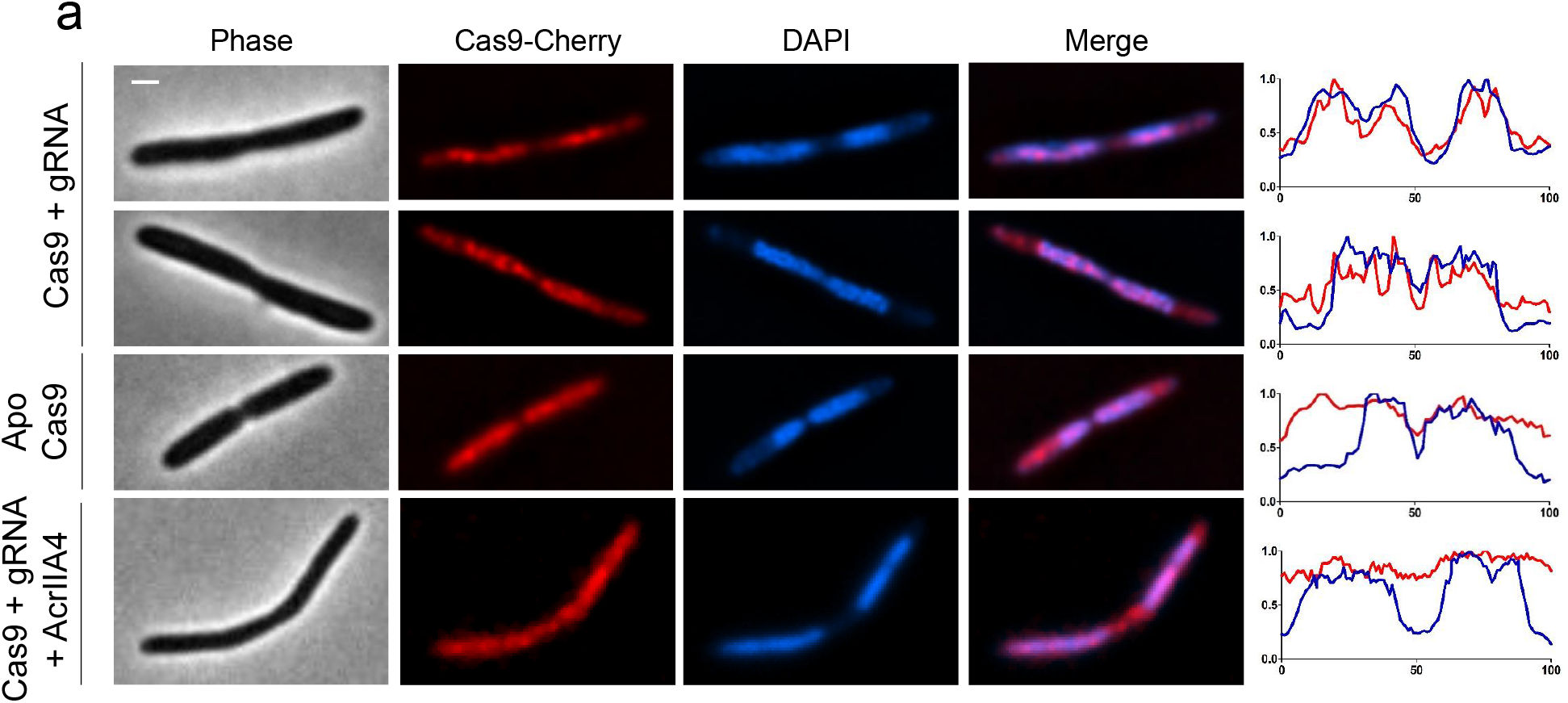
Nucleoid localization is conserved for SpyCas9. Fluorescence microscopy of nucleoid compacted *P. aeruginosa* PA01 cells producing Cas9-sgRNA, ApoCas9, or Cas9-sgRNA co-expressed with anti-CRISPR protein AcrIIA4. Fluorescence intensity map plotted against the long cell axis are shown. Scale bar, 1 μm.

## Discussion

In this study, we report on the cell biology and subcellular organization of the *P. aeruginosa* type I-F system in its native host. We show that Csy1 and Csy4, which are used here as markers of the Csy surveillance complex, are enriched in the nucleoid, consistent with studies of Type I-E Cascade (Vink *et al*., 2020), while the Cas3 nuclease-helicase is distributed in the cytoplasm (schematically illustrated in Fig. 6). Nucleoid localization is not an intrinsic property of these proteins when expressed alone or without crRNAs, but is rather mediated by the assembly of the Csy surveillance complex. This is especially surprising regarding Csy1, as it and other Cas8 family members have been shown to bind DNA non-specifically (Jore *et al*., 2011; Sashital *et al*., 2012; Dillard *et al*., 2018; Vink *et al*., 2020). These observations indicate that the surveillance complex spends most of its time scanning for a match, not in association with the Cas3 nuclease, perhaps avoiding potential detrimental effects. To the best of our knowledge, this is the first study to examine the *in vivo* subcellular organization of Cas3, a universally conserved nuclease in Type I systems. Our observations reveal a new layer of post-translational spatiotemporal regulation in CRISPR-Cas. However, at first glance, the subcellular organization of CRISPR-Cas seems like an inferior approach compared to Type I R-M complexes which localize in the inner membrane (Holubová *et al*., 2000; Holubova *et al*., 2004), likely minimizing host collateral damage and maximizing phage detection. Given this, it remains to be seen whether Csy/Cascade/Cas9 surveillance of the genome is adaptive, as opposed to simply being a by-product of the basic PAM surveillance mechanisms of CRISPR-Cas systems, “forcing” these systems to localize in the genome.

**Figure 6.**
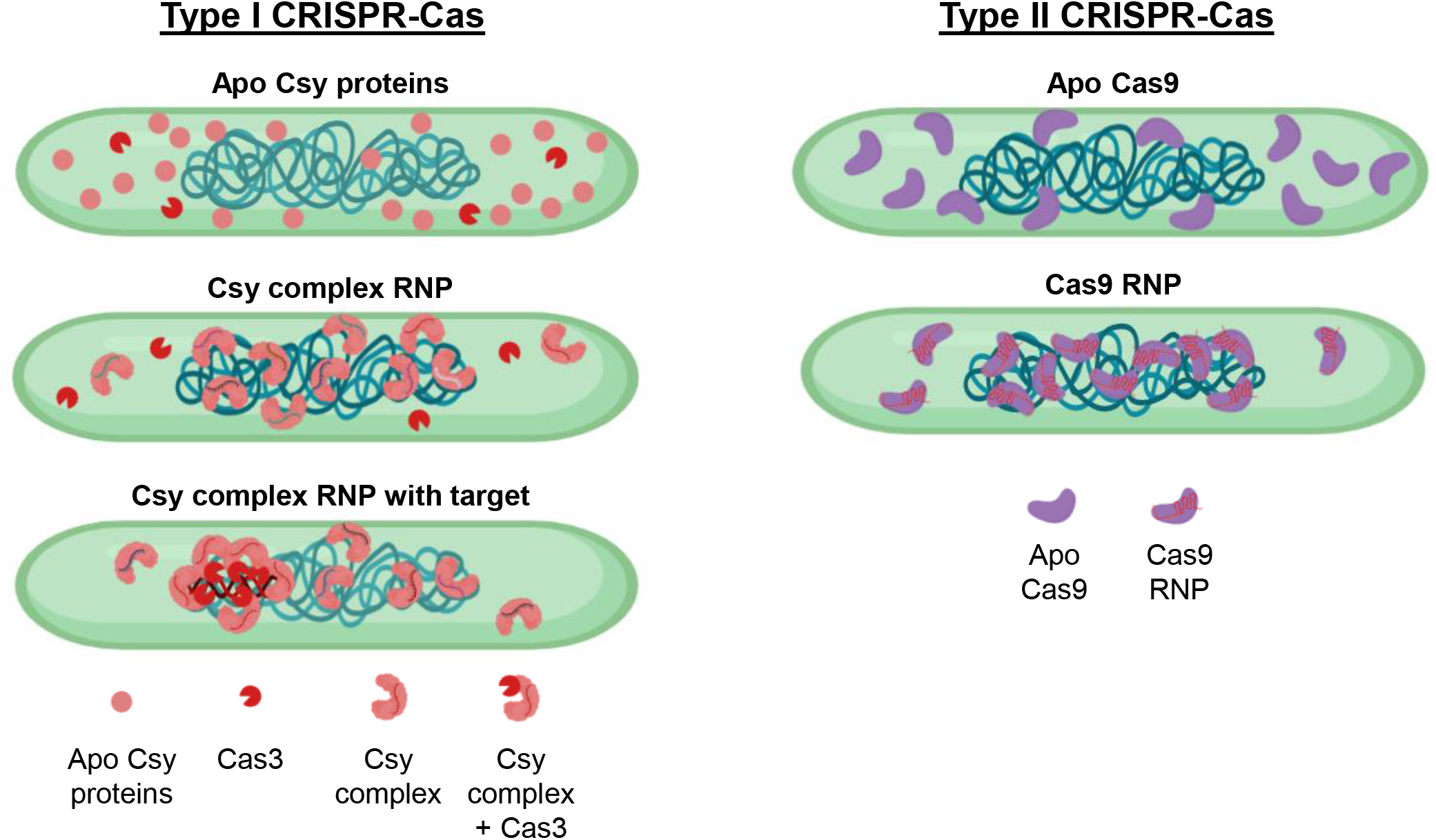
A model for subcellular organization of Type I and Type II CRISPR-Cas systems in bacteria. The subcellular localization of CRISPR-Cas systems are presented for Type I (left panel) and Type II (right panel) systems.

Our observation of differential localization of the Csy complex and Cas3 might present a regulatory strategy to prevent autoimmunity in type I systems, in a way that is not achievable for Class 2 systems (e.g. Cas9 and Cas12). Type II-A Cas9 of *S. pyogenes*, which mediates target recognition as well as cleavage as a single multidomain protein (Jiang and Doudna, 2017), is also enriched in the nucleoid, only when bound to a crRNA. In contrast to the Cas3 nuclease, localization of Cas9 in the nucleoid could pose autoimmunity risks. In fact, a recent study showed that uncontrolled induction Cas9 in *S. pyogenes*, which occurs in the absence of a natural single guide RNA, results in increased self-targeting and auto immunity (Workman *et al*., 2020). Anecdotally, we have also observed that activity of *Listeria monocytogenes* Cas9 is very low, at least in laboratory conditions (Rauch *et al*., 2017). It is possible that because of the single protein mediating DNA-binding and cleavage, that the risks associated with Cas9-mediated toxicity could explain the observed bias towards type I systems in nature.

Our efforts to pin down the target recognition step that mediates nucleoid localization of the Csy complex as well as Cas9 were supported by experiments using Acr proteins, such as AcrIF2, AcrIF11, and AcrIIA4, which specifically block PAM-binding, while AcrIF1 and AcrIF3 act downstream. Perhaps most notably, the inability of AcrIF1 to prevent nucleoid localization, together with observations that crRNA sequence is not important, suggests that PAM-binding is important. Some base pairing in the seed region could contribute to binding strength (Cooper *et al*., 2018), but we do not consider it necessary for nucleoid surveillance due to the abundance of PAM sequences in the genome (*P. aeruginosa* PA14 has 1.2 x 10^6^ 5’-GG-3’ PAM sites). In contrast to our observation, Cascade-DNA interactions were suggested to be mediated by PAM-dependent as well as PAM-independent interactions (Vink *et al*., 2020) because a mutant Cas8 (Csy1 in Type I-F) that is incapable of recognizing PAM decreased, but not completely abolished, its nucleoid localization. In our case, Csy1 nucleoid localization was not observed either in the presence of AcrIF2 or in the absence of crRNA or when Csy1 was expressed alone. Given the divergence of the Cas8 superfamily, it is possible that the mechanistic differences between type I-E and type I-F systems exist.

Lastly, while phage infection has been implicated in upregulation of CRISPR-Cas enzymes in some bacteria and archaea (Agari *et al*., 2010; Young *et al*., 2012; Quax *et al*., 2013), we did not observe significant upregulation or re-localization of the Csy complex or Cas3 during phage infection. Intriguingly, JBD18 could promote clustering of Csy complex and Cas3 as a prophage but not as an infecting phage. Notably, the fluorescent foci formed by dCas3 were quite discrete, suggesting that most dCas3 molecules within the cell are recruited, an observation we found surprising. Whether this truly reflects Cas3 recruitment events or is observed due to catalytic inactivation is worth further investigation. One possible reason for the absence of any Csy/Cas3 re-localization during lytic infection could be the transient nature of the interaction, with a relatively dynamic target like injected phage DNA (Shao *et al*., 2015). Additionally, Csy complexes are loaded with 35 distinct crRNAs, likely concealing relocation of a minority of molecules, despite Csy and Cas3 re-localization being observable for a prophage. Dynamic relocation of molecules near the inner membrane during infection may require investigation using more sensitive imaging techniques including single molecule imaging and total internal reflection fluorescence (TIRF) microscopy (Shashkova and Leake, 2017).

What is the physiological function of the nucleoid enriched CRISPR-Cas enzymes? Several findings have suggested that CRISPR-Cas enzymes participate in cellular functions other than immunity. These include DNA repair, gene regulation, sporulation, genome evolution and stress response (Weiss and Sampson, 2014; Ratner *et al*., 2015; Hille and Charpentier, 2016), however there is no strong evidence for these functions with the Csy complex in *P. aeurginosa*. The molecular basis for most of these alternative functions of Cas proteins are not well understood. A recent study showing SpyCas9 can repress its own promoter using a natural single guide opens the possibility that CRISPR-Cas systems can also function as intrinsic transcriptional regulators (Workman *et al*., 2020). It is therefore possible that crRNAs generated from degenerate self-targeting spacers might direct CRISPR-Cas enzymes in regulating host genes. However, binding of the *E. coli* cascade complex to hundreds of off-target sites does not affect gene expression globally (Cooper *et al*., 2018). Alternatively, it could be concluded that the intrinsic PAM-sensing mechanism of DNA detection relegates Cascade, the Csy complex, and Cas9, to survey the genome, even if that function is neither directly adaptive for cellular functions or defense. Given that affirmative PAM recognition is important for function (and avoidance of CRISPR locus targeting), as opposed to exclusion mechanisms (e.g. genome-wide methylation to prevent restriction enzyme binding), host genome surveillance likely reflects this limitation. Future studies examining these possibilities will provide important insights on the functions and evolutionary limitations of CRISPR-Cas systems in bacteria.

## Materials and methods

### Plasmids, phages and growth media

Plasmids, and primer sequences used in this study are listed in Supplemental Tables 1–2. *P. aeruginosa* UCBPP-PA14 (PA14) strains and *Escherichia coli* strains were grown on lysogeny broth (LB) agar or liquid at 37 °C. To maintain the pHERD30T plasmid, the media was supplemented with gentamicin (50 μg ml-1 for *P. aeruginosa* and 30 μg ml-1 for *E. coli*). Phage stocks were prepared as described previously (Borges *et al*., 2018). In brief, 3 ml SM buffer was added to plate lysates of the desired purified phage and incubated at room temperature for 15 min. SM buffer containing phages was collected and 100 μl chloroform was added. This was centrifuged at 10,000g for 5 min and supernatant containing phages was transferred to a storage tube with a screw cap and incubated at 4 °C. Phages used in this study include DMS3, DMS3m, and engineered DMS3m phage with Acrs (Borges *et al*., 2018).

### Construction of plasmids and strains

Plasmids expressing sfCherry alone, sfCherry tagged with Cas3 orCas9 were previously reported(Mendoza *et al*., 2020). Plasmids expressing Csy1–sfCherry and Csy4-sfCherry were constructed by Gibson assembly in pHERD30T plasmid digested with SacI and PstI. These fusions have ggaggcggtggagcc (G-G-G-G-A) linker sequence in between them. sCherry was amplified from SF-pSFFV-sfCherryFL1M3_TagBFP (kindly provide by Bo Huang lab, UCSF). *csy1* and*csy4*sequences were amplified from PA14. Csy1 and Cas3 are tagged at the N-terminus and Csy4 is tagged at the C-terminus.

Endogenous Csy1–sfCherry and Cas3–sfCherry reporters were previously described (Borges *et al*., 2020). Csy4-sfCherry was constructed in a similar way. The sfCherry gene was inserted with csy4 of PA14 via allelic replacement. The recombination vector pMQ30, which contained sfCherry flanked by homology arms matching csy4 was introduced via conjugation. pMQ30–Csy4-sfCherry, which contains the sfCherry sequence flanked by 123 bp upstream and downstream of csy4 stop codon was cloned in the pMQ30 plasmid between HindIII and BamHI sites using Gibson assembly. Both pMQ30–Csy4-sfCherry contain theGGAGGCGGTGGAGCC sequence (encoding GGGGA) as a linker between sfCherry and csy4. The pMQ30–Csy4-sfCherry construct was introduced into PA14 strains of interest via allelic replacement to generate Csy4-sfCherry. Strains containing the appropriate insertion were verified via PCR.

crRNAs suitable for type I-F system were expressed from I-F entry vectors pAB04. Oligonucleotides with repeat-specific overhangs encoding the spacer sequences that does not have sequence homology with PA14 genome are phosphorylated using T4 polynucleotide kinase (PNK) and cloned into the entry vectors using the BbsI sites. Sequences of the spacers are listed in supplementary table 3. crRNAs are expressed without addition of the inducer arabinose.

### Construction of PA14 lysogens

Lysogens were obtained by first spotting phage onto a bacterial lawn, then streaking out surviving colonies from phage spots. These colonies were screened for phage resistance using a cross-streak method and lysogeny was verified by prophage induction. For maintenance of DMS3m engineered lysogens that expresses Acr proteins AcrIF1, AcrIF2, AcrIF3 during NA treatment, an arabinose inducible plasmid expressing C-repressor of DMS3 was maintained.

### Live-cell imaging and image processing

Fluorescence microscopy was carried out as described previously (Mendoza *et al*., 2020). Unless indicated, overnight cultures were diluted 1:10 in fresh LB medium and grown for 3 hours. For compaction of nucleoid, Nalidixic acid (200 μg/ml) was added for 3 hours or chloramphenicol (200 μg/ml) was added for the last 20 minutes. 0.5 ml cells were centrifuged, washed with 1:10 LB diluted with double-distilled water and finally resuspended in 200–500 μl of 1:10 LB. Cell suspensions were placed on 0.85% 1:10 LB agarose pads with uncoated coverslips. For DNA staining, DAPI (2μg/ml) was added to the cell suspension for 10 minutes and washed twice with 1:10 LB and finally resuspended. A Nikon Ti2-E inverted microscope equipped with the Perfect Focus System (PFS) and a Photometrics Prime 95B 25-mm camera were used for live-cell imaging. Time-lapse imaging was performed using Nikon Eclipse Ti2-E equipped with OKOLAB cage incubator. Images were processed using NIS Elements AR software.

For measuring of single cell fluorescence, region of interest (ROI) were drawn over the phase contrast images of endogenous sfCherry-tagged reporter strains using NIS Elements AR. After subtracting the background, the sfCherry ROI mean intensity values were obtained. The data was analyzed and presented as a scatterplot using GraphPad prism.

For fluorescence intensity maps, an intensity line was drawn over the phase contrast images along the middle long cell axis. Fluorescence values of sfCherry and DAPI along the long axis were extracted. The data was analyzed and presented as a line plot using GraphPad prism.

---

**Supplementary Table - 1.**

| Plasmid name | Information | Reference |
| --- | --- | --- |
| pSG30T-sfCherry-csy1(IF) | pHERD30T with Csy1-sfCherry | This study |
| pSG30T-csy4-sfCherry (IF) | pHERD30T with Csy4-sfCherry | This study |
| pMQ30-csy4-sfCherry |  | This study |
| pSG30T-GpT-mNeonGreen | pHERD30T with capsid GpT-mNeonGreen | This study |
| pHerd30T-C-rep | pHERD30T with DMS3 C-repressor protein | (Borges *et al.*, 2018) |
| pAB014 | I-F entry vectors | (Mendoza *et al.*, 2019) |
| pHerd30T-AcrIF1 | Expresses AcrIF1 | (Bondy-Denomy *et al.*, 2013) |
| pHerd30T-AcrIF2 | Expresses AcrIF2 | (Bondy-Denomy *et al.*, 2013) |
| pHerd30T-AcrIF3 | Expresses AcrIF3 | (Bondy-Denomy *et al.*, 2013) |
| pESN28 | Expresses cherry-Cas9 | (Mendoza *et al.*, 2020) |

**Supplementary Table - 2.**
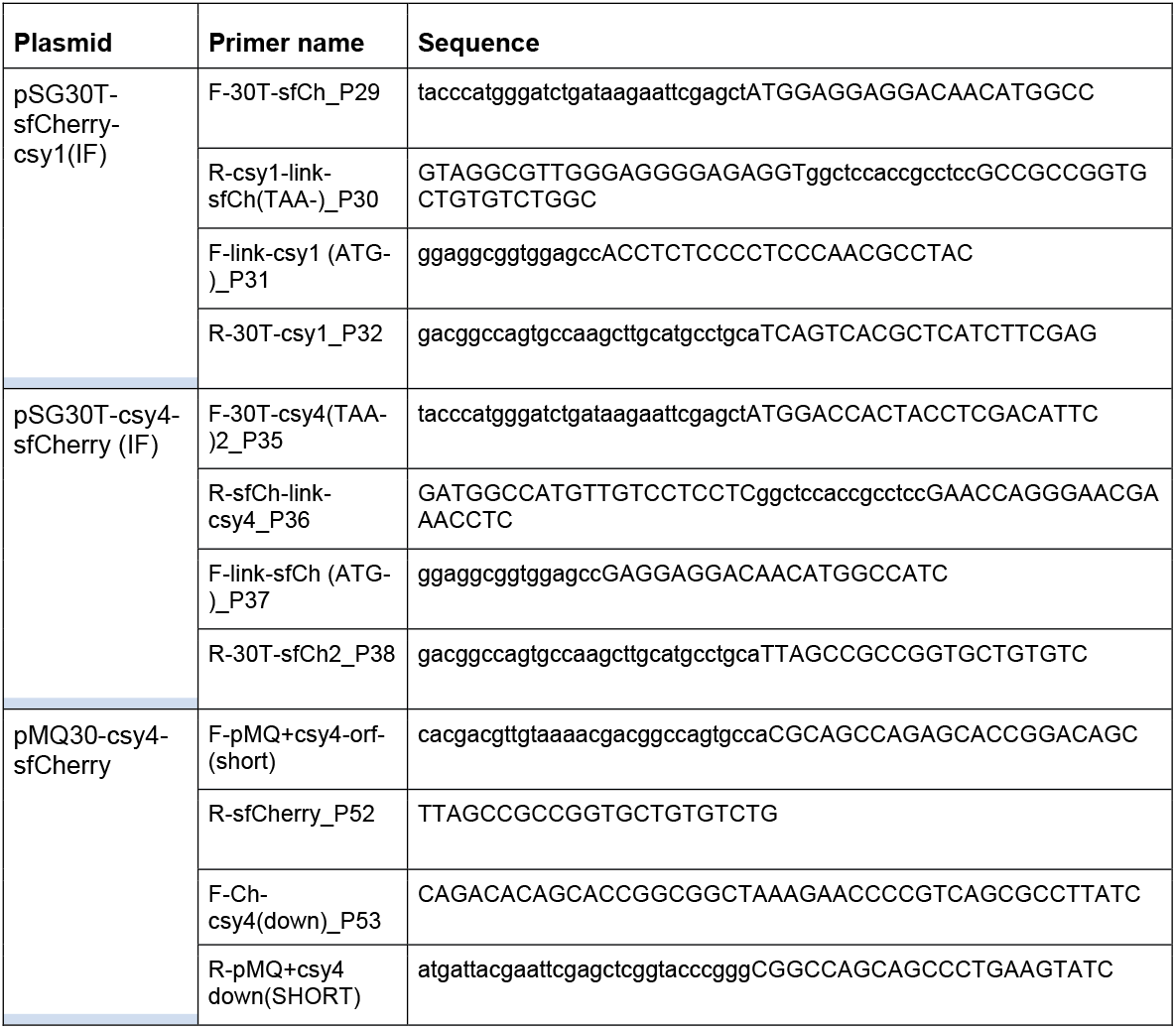
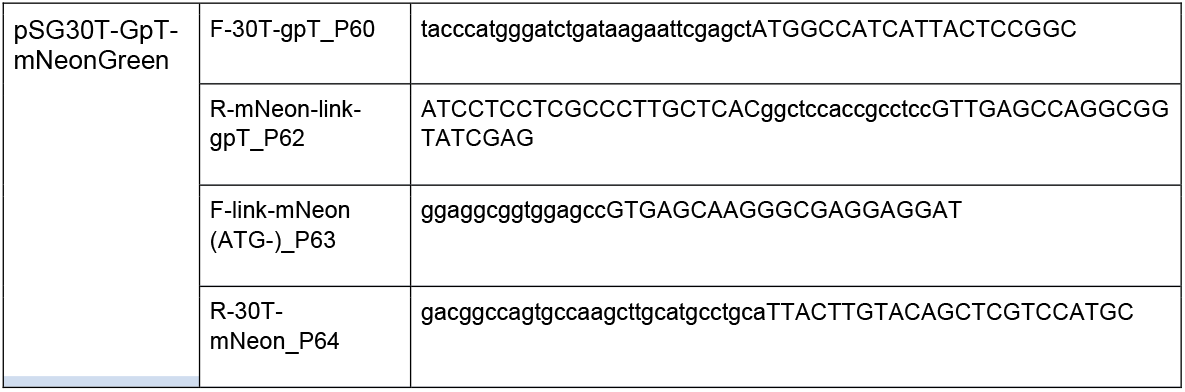

**Supplementary Table - 3.**
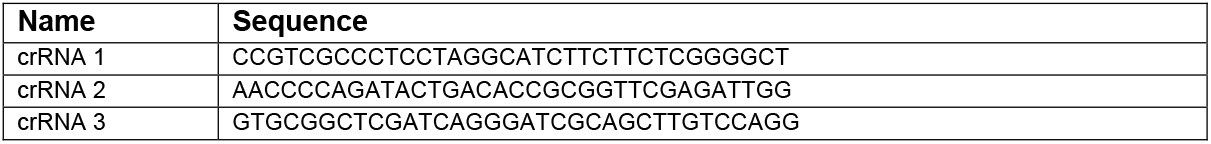

## Conflict of interest

J.B.-D. is a scientific advisory board member of SNIPR Biome and Excision Biotherapeutics and a scientific advisory board member and co-founder of Acrigen Biosciences.

## Acknowledgements

We thank members of the Bondy-Denomy lab for helpful discussions. S.G appreciate helpful discussion and support with Shweta Karambelkar and Bálint Csörgő. This project in the Bondy-Denomy lab was supported by the UCSF Program for Breakthrough Biomedical Research funded in part by the Sandler Foundation and an NIH Director’s Early Independence Award DP5-OD021344.

---

**Figure S1.**
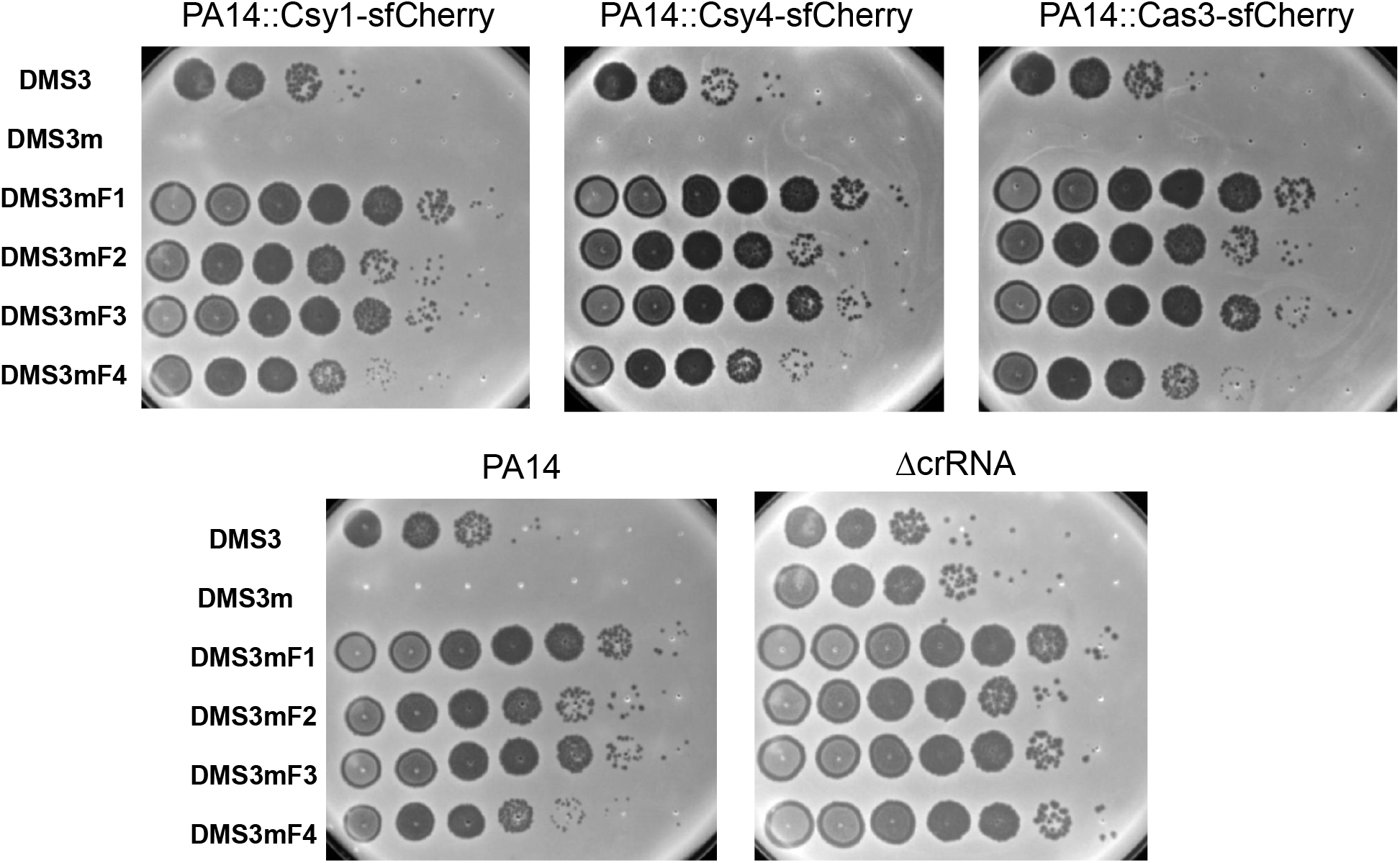
sfCherry-tagged endogenous type I-F reporters are functional. A spot titration of DMS3, DMS3m and DMS3m engineered phages that expresses Acr proteins AcrIF1, AcrIF2, AcrIF3 or AcrIF4 on sfCherry-tagged, WT PA14, and crRNA deletion strains.

**Fig. S2.**
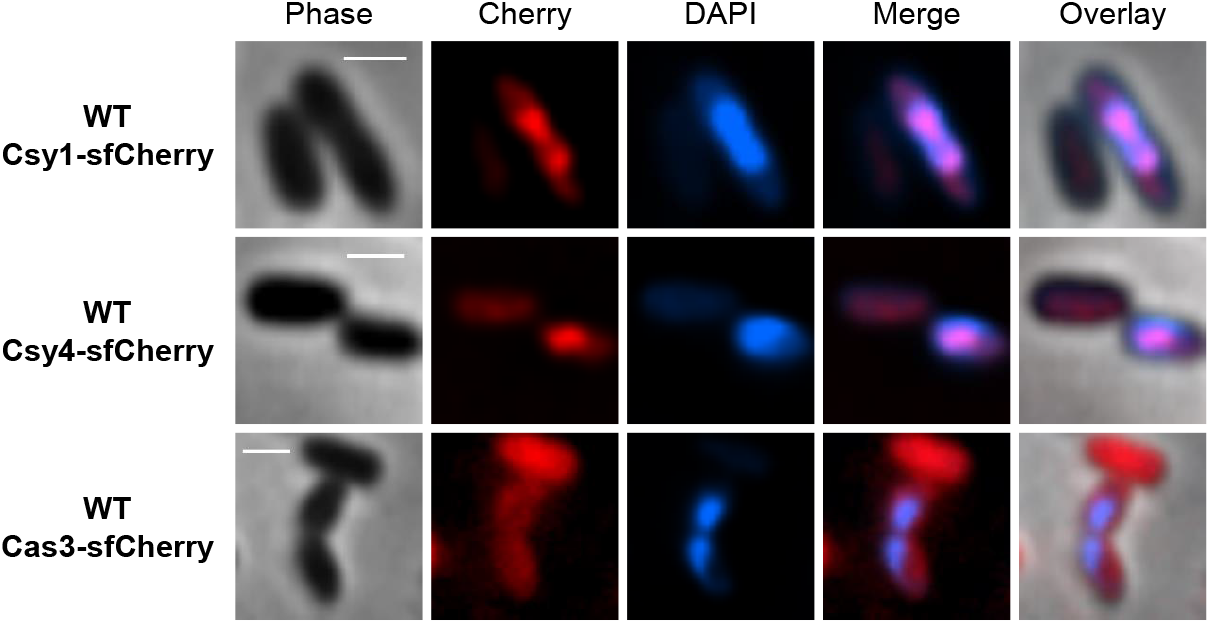
Nucleoid enrichment of Csy complex is not due to DNA damage. Fluorescence microscopy of wild-type cells expressing Csy1-sfCherry, Csy4-sfCherry or Cas3-sfCherry in chloramphenicol-treated nucleoid compacted cells.

**Fig. S3.**
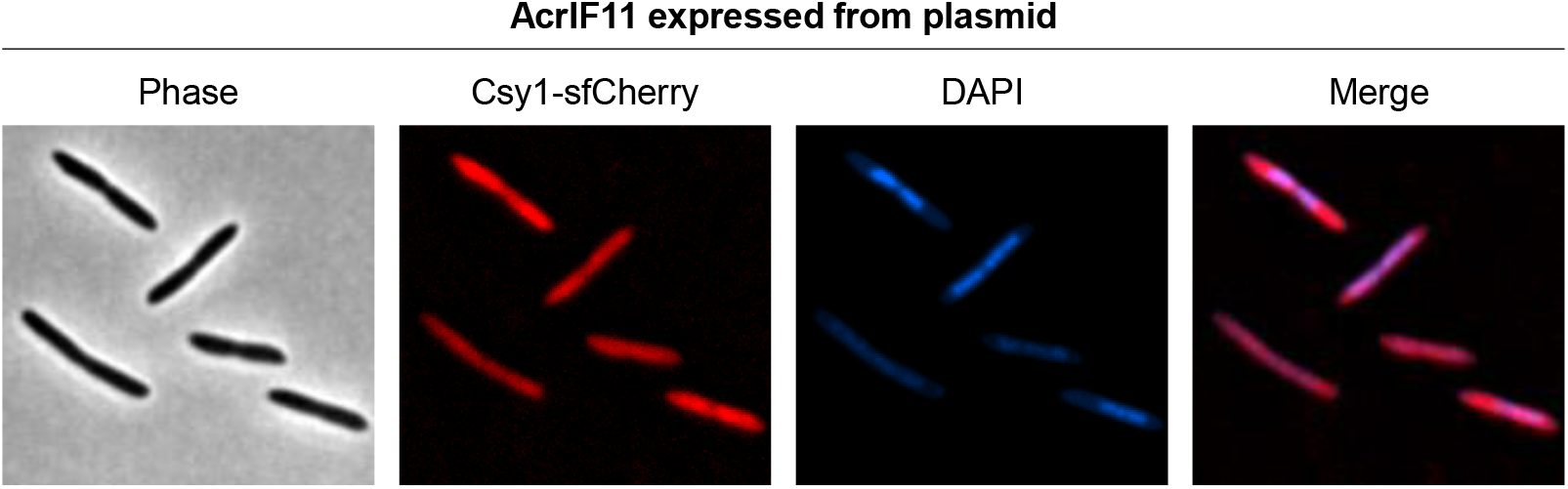
AcrIF11 blocks nucleoid localization of the Csy complex. Fluorescence microscopy of nucleoid compacted wild-type cells producing Csy1-sfCherry and expressing anti-CRISPR proteins AcrIF11 from a plasmid.

**Fig. S4.**
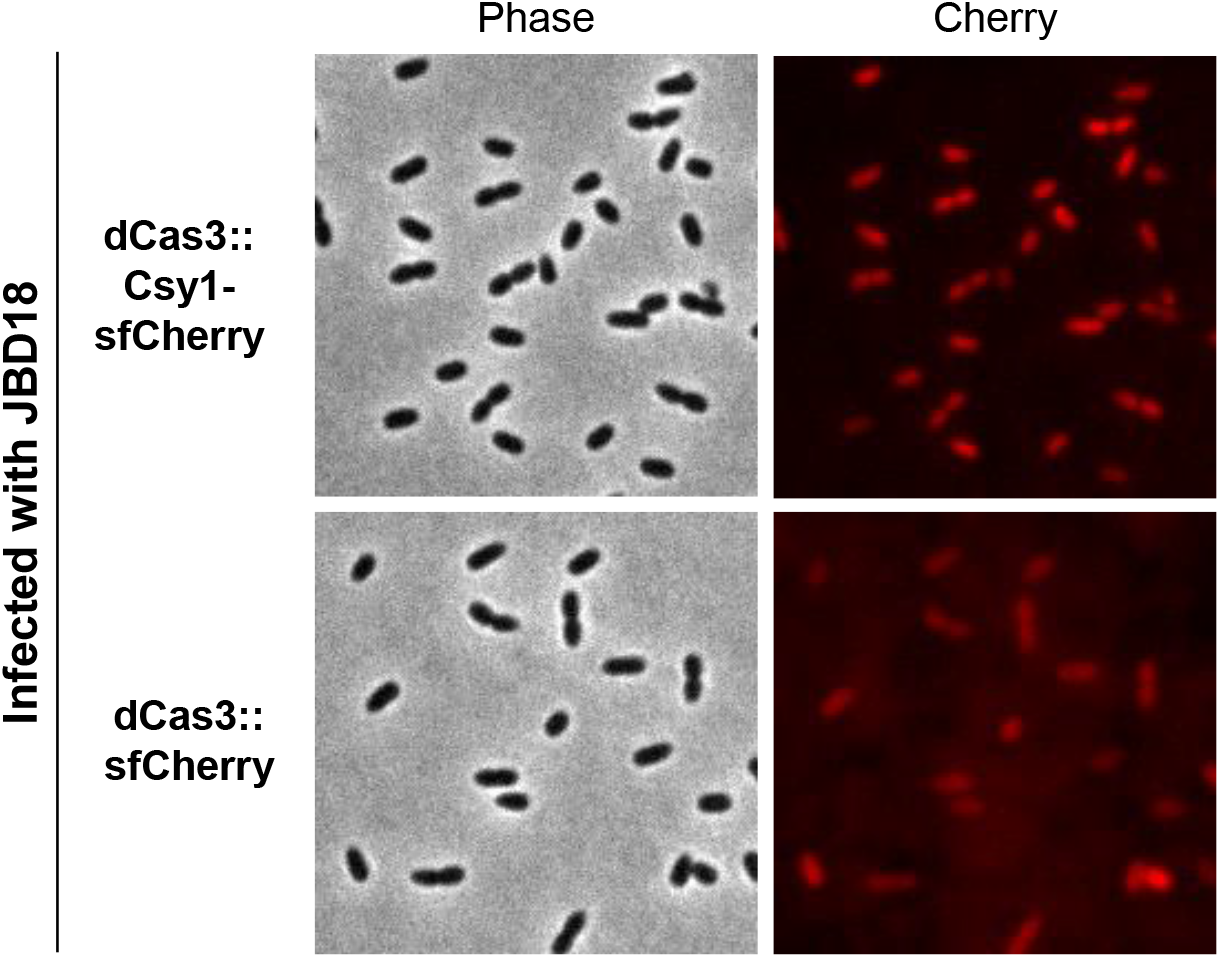
Phage infection does not modulate Csy1 localization. Fluorescence microscopy Csy1-sfCherry (in dCas3 background) or dCas3-sfCherry expressing cells infected with JBD18.

